# Influenza C incidence and herd immunity in Lancaster, UK, in the winter of 2014-2015

**DOI:** 10.1101/061705

**Authors:** Kate V Atkinson, Lisa A Bishop, Glenn Rhodes, Nico Salez, Neil R McEwan, Matthew J Hegarty, Julie Robey, Nicola Harding, Simon Wetherell, Robert M Lauder, Roger W Pickup, Mark Wilkinson, Derek Gatherer

## Abstract

Influenza C is not included in the annual seasonal influenza vaccine, and has historically been regarded as a minor respiratory pathogen. However, recent work has highlighted its potential role as a cause of pneumonia in infants. We performed nasopharyngeal or nasal swabbing and/or serum sampling (n=148) in Lancaster, UK, over the winter of 2014-2015. Using enzyme-linked immunosorbent assay (ELISA), we estimated a seropositivity of 77%. By contrast, only 2 individuals, both asymptomatic adults, were influenza C-positive by polymerase chain reaction (PCR). Deep sequencing of nasopharyngeal samples produced partial sequences for 4 genome segments in one of these patients. Bayesian phylogenetic analysis demonstrated that the influenza C genome from this individual is evolutionarily distant to those sampled in recent years and represents a novel genome constellation, indicating that it is a product of a decades-old reassortment event. Although we find no evidence that influenza C was a significant respiratory pathogen during the winter of 2014-2015 in Lancaster, we confirm previous observations of seropositivity in the majority of the population. We calculate that this level of herd immunity would be sufficient to suppress epidemics of influenza C and restricts the virus to sporadic endemic spread.

## Introduction

### Clinical presentation

Influenza C (family *Orthomyxoviridae*, genus *Influenzavirus* C, species *Influenza* C *virus*) produces malaise and coryza when administered to susceptible healthy adult volunteers, with fever in a minority of cases [Joosting et al., 1968]. Historically, influenza C has been regarded as the least serious of the three species of influenza infecting humans, and seasonal vaccination programmes have been confined to influenzas A and B. More recent studies in Finnish army recruits confirmed influenza C’s production of a mild respiratory illness in healthy adults, with only occasional complications [Kauppila et al., 2013].

However, in a paediatric context, acute respiratory illness and/or pneumonia have been reported as a consequence of influenza C infection [Calvo et al., 2006; Matsuzaki et al., 2007; Matsuzaki et al., 2003; Moriuchi et al., 1991; Peng et al., 1996; Principi et al., 2012; Shimizu et al., 2015] especially in those under 2 years old [Matsuzaki et al., 2006], as well as vomiting, diarrhoea, acute otitis media [Laxdal et al., 1966], a high rate of hospitalization [Gouarin et al., 2008] and even acute encephalopathy [Takayanagi et al., 2009]. There is increased recognition that under-reporting of influenza C in children is a problem [Pabbaraju et al., 2013]. This growing awareness of the paediatric clinical importance of influenza C raises the issue of its inclusion in the annual seasonal influenza vaccine, or its position as a candidate for vaccine development specifically for infants.

### Epidemiology

Nearly 40% of adult volunteers were susceptible to administered influenza C [Joosting et al., 1968]. The 60% who did not develop disease after experimental exposure is neatly consonant with observation of seropositivity levels of 59% in Spain [Manuguerra et al., 1994], 61% in France [Manuguerra et al., 1992] and 57% in Brazil [Motta et al., 2000], and suggests that seropositivity may possibly confer resistance. By contrast, other studies have suggested that antibodies against influenza C tend to be more universal: 100% in an isolated Philippine village [Nishimura et al., 1987] and in US adults and children [Hilleman et al., 1953], 90% in Czechoslovakia, 86% in the Soviet Union [Vasil'eva et al., 1985] and 70% in East Germany [Tumova et al., 1983]. Antibody titre levels among those classed as seropositive, varied widely. Some studies have also found age-structured variability: in California, seropositivity of 64% in children under 5 but 98% in adults [Dykes et al., 1980]; in Japan, 40-50% in early childhood to nearly 100% in adulthood [Kaji et al., 1983]; in Louisiana, 47% in children to 96% in younger adults, but then a decline to 18% in the over-65s [O'Callaghan et al., 1980]; in France, 46% seropositivity in children, 76% in younger adults, but only 44% in the over-50s [Manuguerra et al., 1992].

Influenza C does not appear to be seasonal, based on contemporaneous two-year surveys of its occurrence in Bucharest and Japan from 1988-1990 [Ionita et al., 1992; Moriuchi et al., 1991]. Using this observation together with the seropositivity data, it is possible to propose several epidemiological scenarios. The first of these is that influenza C is essentially an endemic virus in human populations, with more or less lifelong immunity conferred by exposure but, in the absence of a vaccination programme, a sufficient supply of newborns and unexposed adults to provide a host population. The decline in seropositivity in later life, found in at least two studies [Manuguerra et al., 1992; O'Callaghan et al., 1980], potentially due to immunosenescence, would then provide the virus with opportunities to infect individuals for a second time. The second scenario is that the virus is only intermittently epidemic. The variation in seropositivity according to place, time and individual age is therefore a reflection of previous epidemic history in different locations. The third scenario is that the virus is endemic but antigenically variable over time. Seropositivity would therefore be an unreliable guide to the true immune status of any individual. Individuals may acquire immunity, but this will eventually disappear as its value is eroded by antigenic drift, for which there is some evidence in influenza C [Chakraverty, 1978].

### Phylogenetics and molecular evolution

The rate of nucleotide substitution is lower in influenza C than in A and B [Buonagurio et al., 1986; Gatherer, 2010; Muraki et al., 1996; Yamashita et al., 1988]. Like the other influenza viruses, influenza C has a segmented RNA genome, and reassortment has been detected [Buonagurio et al., 1986; Gatherer, 2010; Matsuzaki et al., 2003; Moriuchi et al., 1991; Peng et al., 1994; Racaniello and Palese, 1979]. There is also evidence of positive selection for evasion of the host immune system at two residues in the receptor-binding domain of the haemagglutinin-esterase (HE) protein, but the overall ratio of non-synonymous to synonymous substitutions (omega) across the genome is low, individual proteins ranging from 0.05 to 0.13 [Gatherer, 2010]. The low levels of omega indicate a virus that is well adapted to its host, but the presence of positive selection in the HE receptor-binding domain also indicates selective pressure from the host immune system. This provides a molecular explanation for the observed antigenic drift [Chakraverty, 1978] and some evidence against the scenario that humans are likely to acquire lifelong immunity. Influenza C would therefore resemble influenza A and B in that a new vaccine would be required every time the antigenic drift had reached a certain extent, potentially annually.

The issue of endemicity versus sporadic epidemics also remains unresolved. Only one candidate epidemic surge has been identified, in Japan in 2004 [Matsuzaki et al., 2007]. The existence of reassorted strains indicates that double infection with two or more strains cannot be very infrequent, implying that it ought to be possible to detect numerous (or at least >1) strains co-circulating both temporally and geographically, previously demonstrated in Japan [Matsuzaki et al., 2007]. Indeed, a continually shifting pattern of segment combinations, referred to as genome constellations [Gatherer, 2010], is observed when full genomes are studied, a phenomenon also seen in influenza B [Chen and Holmes, 2008]. Eight genome constellations circulating in the 1990s differed from the genome constellations present in a set of reference genomes from the 1940s to the 1980s [Gatherer, 2010].

## Methods

### Patient recruitment

Lancaster (54.05°N 2.80°W) is a small city with a population of 45,000 rising to 141,000 when surrounding towns and villages are included. The permanent resident population is >95% white and 18% are over age 65. 3 cohorts of participants were approached: 1) staff and students at Lancaster University, 2) patients attending a general practitioner (GP) consultation, 3) patients attending hospital clinics. After informed consent was given, patients with coryza and/or other symptoms consistent with respiratory infection, were classified as the symptomatic group (*n*=71) and the remainder as asymptomatic (*n*=77). Nasopharyngeal (or nasal) swabbing, blood sampling, or both, were performed on the patients, according to consent. Sample collection was performed from November 2014 to May 2015.

### Sample processing

Nasopharyngeal swabs (MW951SENT, Medical Wire) were used to remove cells and mucus from the rear wall of the nasopharynx or nose (according to consent) of patients, and the tips then snapped off directly into Sigma Virocult^®^ medium.

Blood was drawn from forearm veins into Beckton Dickinson Vacutainer^®^ tubes containing clot activator or from a finger prick, according to consent, using Beckton Dickinson Serum Separator^®^ tubes (SST™). Serum was separated at 1000-2000g for 10 minutes (for arm samples) or at 6000-15000g for 90s (for finger-prick samples) and then stored at -80°C.

RNA was extracted from the nasopharyngeal swabs using a MagMAX™ Viral RNA Isolation Kit (Ambion). The quality and quantity of RNA extracted from samples was assessed by spectrophotometry using the NanoDrop^®^ 1000 Spectrophotometer V3.3.0 (Thermo Fisher Scientific). cDNA was prepared using a High-Capacity RNA-to-cDNA™ Kit (Applied Biosystems^®^, Life Technologies™) and a Veriti^®^ Thermal Cycler (Applied Biosystems^®^, Life Technologies™). The samples were incubated at 37°C for 60 minutes, before stopping the reaction at 95°C for 5 minutes and then holding at 4°C. Once completed, the plates were stored at -20°C.

Polymerase chain reaction (PCR) was then performed using a 7500 FAST Real-Time PCR system (Applied Biosystems^®^, Life Technologies™) with thermo-cycling carried out as follows: one cycle of 95°C for 10 min and 45 cycles of 95°C for 15 s and 60°C for 1 min. PCR primers for influenza C were as used previously [Salez et al., 2014]. Samples judged positive after quantitative PCR were processed using the Illumina Nextera XT library kit and deep sequenced in 2x126bp format using an Illumina HiSeq2500 system.

Enzyme-linked immunosorbent assay (ELISA) was performed on the serum samples using influenza C antigen as previously described [Salez et al., 2014] and goat anti-human HRP-conjugated secondary antibody (ab6858, Abcam^®^) with SureBlue™ TMB Microwell Peroxidase Substrate solution. Absorbance was measured at 450nM using a Wallac Victor2™ (Perkin Elmer) plate reader. Anti-influenza C IgG was quantified by calibration of the peroxidase reaction against a standard dilution series of IgG concentrations. The threshold for seropositivity was placed at 2 standard deviations above the mean level of the negative control serum.

### Genome segment sequence assembly

Illumina reads were trimmed of adapters and other non-genomic elements using CutAdapt 1.1 [Martin, 2011: https://pypi.python.org/pypi/cutadapt], fastq-mcf 0.11.3 [Aronesty, 2013: https://expressionanalysis.github.io/ea-utils], and trim_galore(http://www.bioinformatics.babraham.ac.uk/projects/trim_galore/), within the Read_cleaner pipeline (Gatherer, unpublished, see Supplementary Data Pack). Ethical approval required that no genetic material remain within the samples which could enable identification of patients. Therefore, human genome and transcriptome sequences were removed by iterative alignment onto the NCBI, Ensembl and UCSC human iGenomes (http://support.illumina.com/sequencing/sequencing_software/igenome.html), first using bowtie 1.1.1 [Langmead et al., 2009: http://bowtie-bio.sourceforge.net/index.shtml], then BWA 0.7.12-r1039 [Li and Durbin, 2010: http://bio-bwa.sourceforge.net] within the Valet pipeline (Gatherer, unpublished, see Supplementary Data Pack). Following each alignment, extraction of unaligned reads was achieved using samtools 0.1.19 [Li et al., 2009: http://samtools.sourceforge.net/] and the next alignment commenced. Bowtie, BWA and samtools were co-ordinated using the Vanator pipeline [Jarrett et al., 2013: https://sourceforge.net/projects/vanator-cvr]. The resulting trimmed and cleaned reads are available from the Sequence Read Archive (https://www.ncbi.nlm.nih.gov/sra/: BioSamples SAMN05954290 and SAMN05954291, Runs SRR4733498 and SRR4733494)

Influenza C genome C/Victoria/2/2012 (Genbank ref. KM504282) was selected as a representative of recently circulating influenza C and alignment of cleaned reads carried out using bowtie within the Valet pipeline. Consensus sequences were constructed using samtools 0.1.19 (bcftools and vcfutils functions). C/Victoria/2/2012 was used to fill gaps in the consensi and the bowtie alignment repeated. This cycle was performed until a stable consensus was obtained for each genome segment. The same process was repeated using BWA and combined consensi obtained. Alignment of reads to the final consensi was examined with Tablet [Milne et al., 2013: https://ics.hutton.ac.uk/tablet]. Resulting assemblies of more than 200 bases were submitted to GenBank (references KY075640 - KY075642). The remaining smaller fragments, along with composite partial segments used in phylogenetic analysis, are available in the Supplementary Data Pack. The new strain of influenza C identified was designated C/Lancaster/1/2015.

### Phylogenetics and genome constellations

Sequence alignments of composite partial segments with full influenza C genomes from GenBank, were performed using Muscle [Edgar, 2004] in MEGA [Kumar et al., 2008: http://www.megasoftware.net] and neighbour joining trees [Saitou and Nei, 1987] constructed. Clock-like behaviour in sequence evolution on those trees was checked using TempEst [Rambaut et al., 2016: http://tree.bio.ed.ac.uk/software/tempest]. Bayesian phylogenetic analysis was performed in BEAST v.1.8.3 [Drummond and Rambaut, 2007: http://tree.bio.ed.ac.uk/software/beast/]. A Tamura [1992] 3-parameter (T93+G) substitution model, coalescent constant size tree prior and relaxed lognormal clock were run for 100 million iterations in BEAST, as previously [Gatherer, 2010]. A burn-in of 25% of all trees was used to create the consensus tree. Genome constellations were determined by establishing the clade, as defined by Gatherer [2010], in which each genome segment was located.

## Results

### Participants

Of the 148 participants, 69 were male and 79 female. 71 were symptomatic and 77 asymptomatic. Distribution of male and female participants within symptomatic and asymptomatic groups was assessed by a 2x2 chi-square test and was not statistically significant. Except for a relative excess of age group 20-29 participants (mostly from the university), age approximated a normal distribution. The summary clinical presentation within the symptomatic group, graph of age distribution and details of the chi-square tests are available in the Supplementary Data Pack.

### Influenza C seropositivity

Of the 148 participants, 129 consented to donate serum. Of these 99 were seropositive and 30 negative, giving a figure of 77% seropositivity. Figure 1 shows the anti-influenza C IgG concentration by age. Gender differences in seropositivity were also nearly absent (male 2.5mg/dl, female 2.3mg/dl) with no statistical significance on t-test, but symptomatic individuals had slightly more IgG (symptomatic 2.6mg/dl, asymptomatic 2.2mg/dl), significant on a t-test at p<0.05. A Mann Whitney U-test was performed on the distribution of seropositive individuals between each age group, and was not statistically significant (see Supplementary Data Pack).

**Figure 1:**
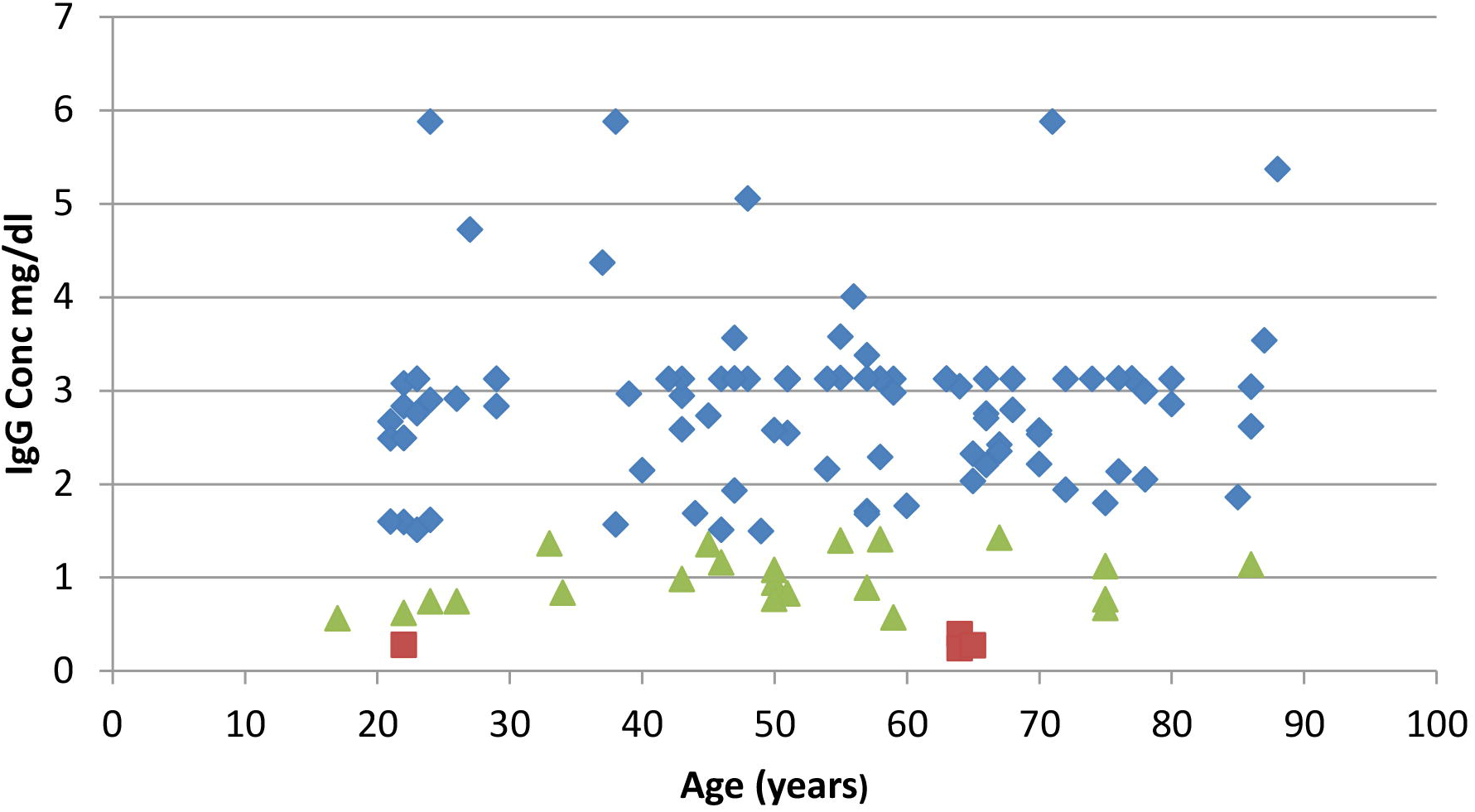
Anti-influenza C IgG concentration. (mg/dl), plotted for each individual against age. Blue: <2 standard deviations above negative control; green: 1-2 standard deviations above negative control; red: <1 standard deviation above negative control.

### Incidence of detectable virus

Two participants out of 148 (1.4%) were detected as positive for influenza C using PCR. Both were asymptomatic. On deep sequencing (SRR4733498 and SRR4733494), only one patient showed sufficient levels of influenza C reads for genome assembly to be attempted (SRR4733498). If the other individual is a false positive by PCR, the population incidence may therefore be 0.7%. This figure is compromised by the fact that the sample is not randomly selected, but is deliberately enriched for symptomatic individuals (71/148, 48%). Since incidence is also often given as positive individuals per symptomatic case, and neither positive individual was symptomatic; on that strict formulation the incidence is 0%. In view of this, we can say little other than that influenza C incidence is low and probably accounted for <1% of coryza and other respiratory disease in Lancaster during the winter of 2014-2015.

### Genetic relationships of isolated influenza C genome segments

Partial genome segment sequences were obtained from deep sequencing for segments 1, 5, 6 and 7, encoding PB2, NP, M1/CM2 and NS1/NS2 respectively. Those greater than 200 bases are deposited in GenBank, accession numbers KY075640 - KY075642 and the remainder are available in the Supplementary Data Pack. Insufficient reads were available to assemble the other segments. Although breadth of coverage across segments is low (ranging from 22% in segment 5 to 32% in segment 6), there is sufficient genetic information to assign each fragment to a clade as defined by Gatherer [2010], using Bayesian phylogenetics. Plotting of the root-to-tip genetic distance on a neighbour-joining tree using TempEst showed that molecular clocks apply best to segments 2 and 7 (PB2 and NS1/NS2), but that both segments 5 and 6 (NP and M1/CM2) have lower root-to-tip distances for C/Lancaster/1/2015 than expected. Figures 2 and 3 shows the TempEst plots for segments 1 and 6 (PB2 and M1/CM2), giving examples of clock-like and non-clock-like behaviour, respectively. The TempEst plots for segments 5 and 7 (NP and NS1/NS2) are Supplementary Figures 3 and 4 respectively.

**Figure 2:**
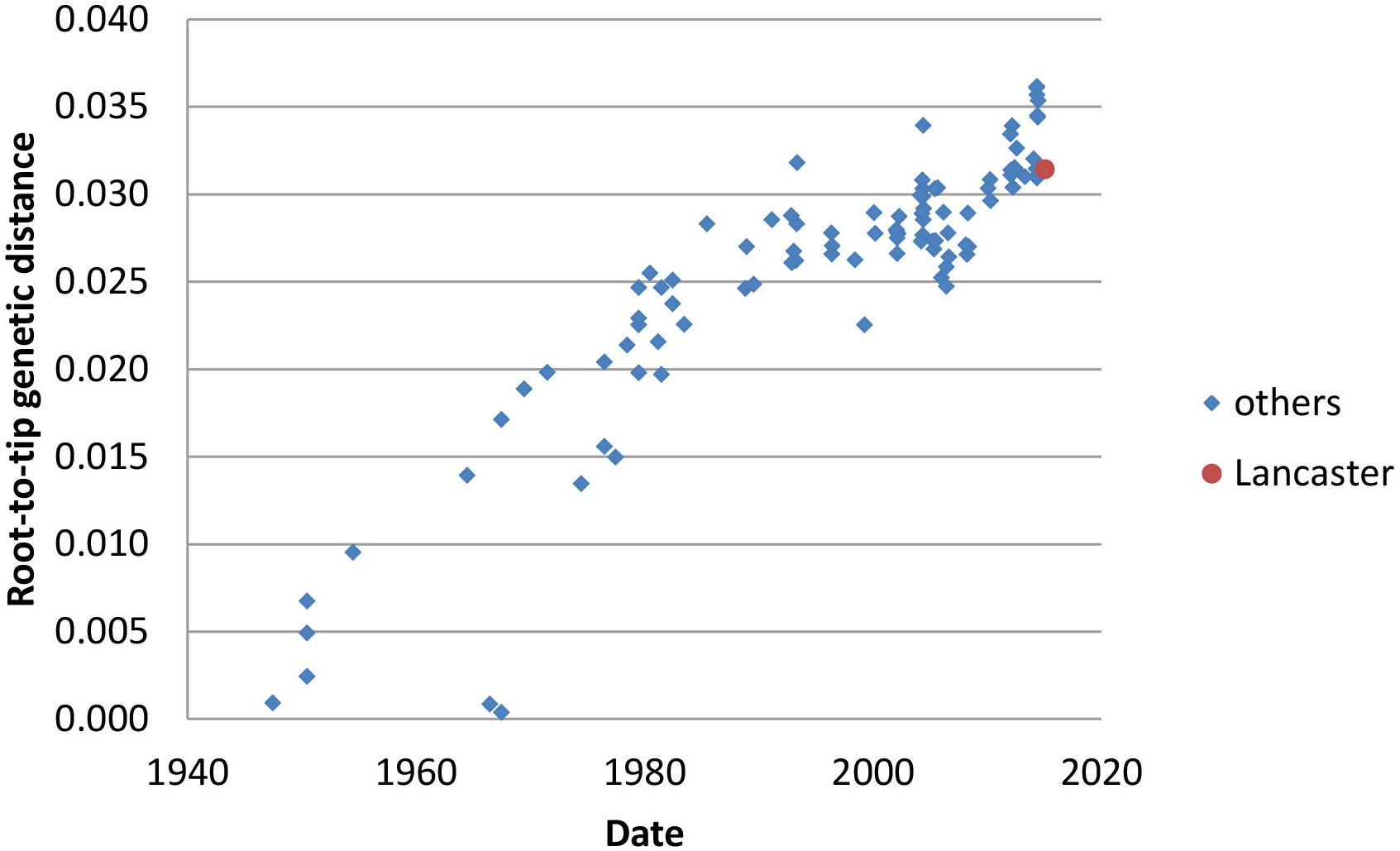
Root-to-tip distance in a neighbour joining tree for segment 1. (encoding PB2) of the influenza C genome. 100 full-length or near full-length genome segments (2365 bases) are used plus the 724 discontinuous bases of segment 1 derived from deep sequencing. C/Lancaster/1/2015 has a degree of divergence from the root consistent with molecular clock-like behaviour in its lineage.

**Figure 3:**
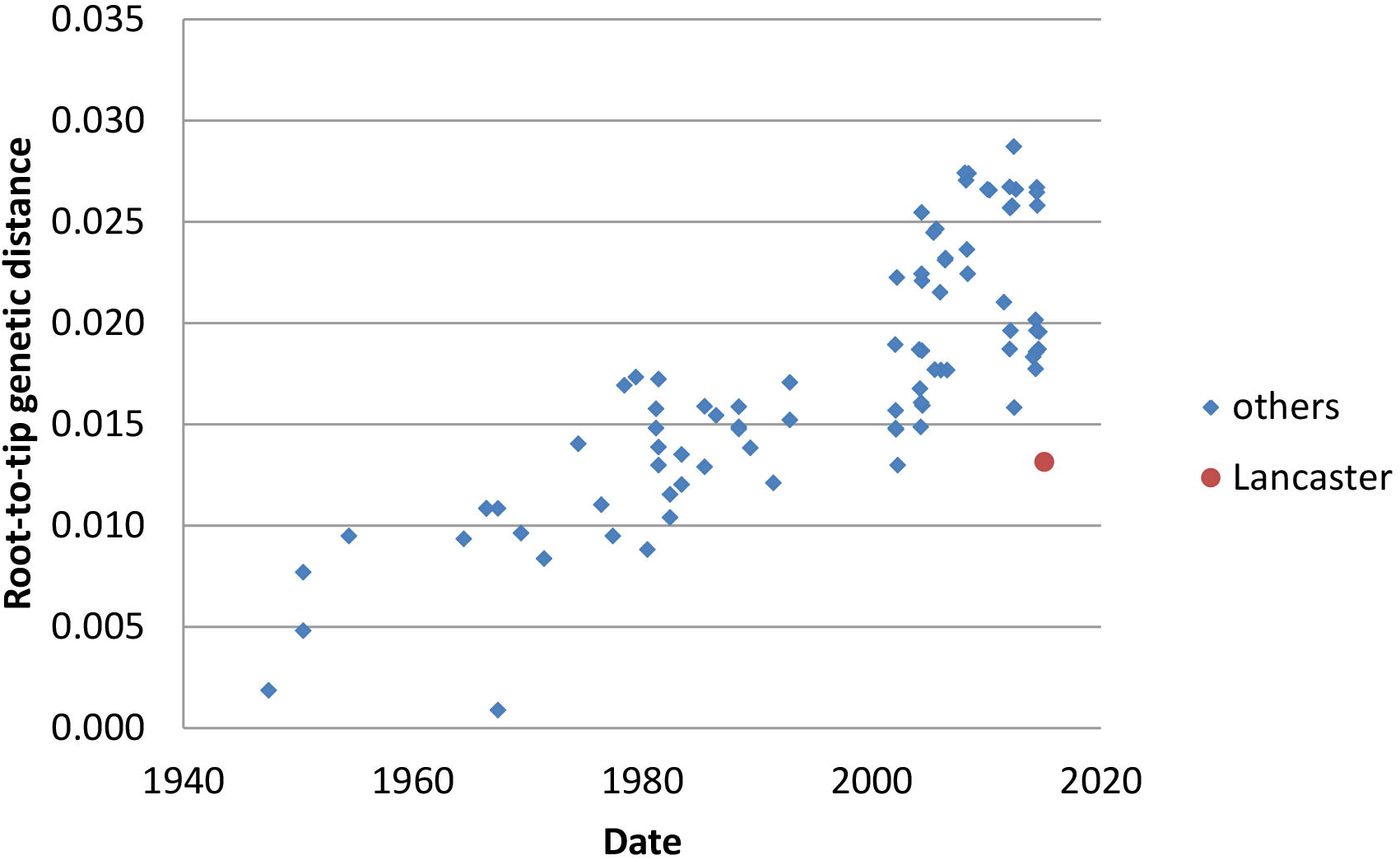
Root-to-tip distance in a neighbour joining tree for segment 6. (encoding M1/CM2) segment of the influenza C genome. 86 full-length or near full-length genome segments (1180 bases) are used plus the 380 discontinuous bases of segment 6 derived from deep sequencing. C/Lancaster/1/2015 is less divergent from the root than it should be given its known sampling date, consistent with a perturbation of molecular clock-like behaviour in its lineage.

Clade memberships were determined by examination of Bayesian phylogenetic trees produced in BEAST, following the classificatory scheme of [Gatherer, 2010] and then annotated onto the neighbour-joining trees used for the molecular clock analysis. Figure 4 shows the tree for segment 5 (encoding NP), demonstrating that C/Lancaster/1/2015 belongs to the C/Miyagi/1/93 clade, and not to the C/Greece/79 and C/pig/Beijing/81 clades circulating in recent isolates. Figure 5 shows the tree for segment 7 (encoding NS1/NS2) has an even more distant relationship to recent genomes, being part of the C/Sapporo/71 clade last seen in 1979. The phylogenetic trees for PB2 and MP are given in Supplementary Figures 1 and 2, and further confirm the genetic distance between C/Lancaster/1/2015 and other recently sequenced genomes. Clade memberships are then synthesised to derive the relationship between C/Lancaster/1/2015 and defined genome constellations (Table 1).

**Figure 4:**
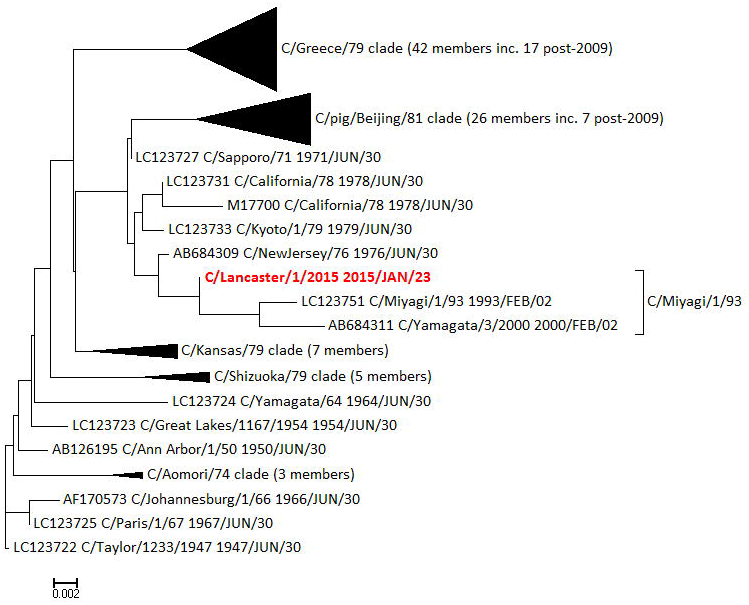
Supplementary Figure 4: Root-to-tip distance in a neighbour joining tree for segment 7. (encoding NS1/NS2) segment of the influenza C genome. 134 full-length or near full-length genome segments (935 bases) are used plus the 288 discontinuous bases of segment 7 derived from deep sequencing. C/Lancaster/1/2015 has a degree of divergence from the root consistent with molecular clock-like behaviour in its lineage.

## Discussion

### Herd immunity to influenza C

Our participant group were 77% seropositive to influenza C. This is slightly higher than the 57-61% levels from studies in western Europe and Brazil [Manuguerra et al., 1992; Manuguerra et al., 1994; Motta et al., 2000], within the range of the 70-90% found in eastern Europe [Tumova et al., 1983; Vasil'eva et al., 1985] but still considerably short of those studies reporting universal seropositivity in the USA and east Asia [Hilleman et al., 1953; Nishimura et al., 1987]. As in previous studies, our antibody titre levels were widely variable among those classed as seropositive, and our choice of threshold is purely statistical. However, we also found no statistically significant age-structured or gender-structured variability in seropositivity (Figure 1). This is at variance with some previous studies in the USA, Japan and Europe [Dykes et al., 1980; Kaji et al., 1983; Manuguerra et al., 1992; O'Callaghan et al., 1980]. It should also be noted that many serological studies on influenza C are now some decades old and techniques have varied over the years, so individual studies are not necessarily directly comparable. We also cannot exclude the possibility of some cross-reactivity of our influenza C antigen with antibodies to other influenza viruses, but this is also an issue in all previous studies.

Whatever the source of the initial antigenic stimulus for the production of anti-influenza C IgG, such seropositivity may be equivalent to immunity to influenza C, even if of a temporary or partial nature, and this may have implications for the epidemiology of the virus. We are not aware of any study on the reproductive number (R_0_) of influenza C, although extensive studies have been performed for influenza A [reviewed by Biggerstaff et al., 2014]. If we assume that the R_0_ of 1.28 for seasonal influenza A, calculated by Biggerstaff et al. [2014] as a median of 47 published values, also applies to influenza C, then the herd immunity threshold (HET) – at which R_t_ would be reduced to 1, and an epidemic would be unsustainable – is:

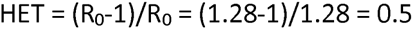

This implies that 50% immunity in the population would be sufficient to suppress any epidemic outbreak of influenza C. Our level of 77% seropositivity may therefore explain why the influenza C virus was so difficult to detect in our participant group. We also note that it is not relevant to this calculation if our 77% partially represents cross-reactive antibodies against influenza A and B. Regardless of the virus type that initially produced the antibodies, their binding to influenza C antigen in ELISA suggests that they may be effectively protective *in vivo* and would contribute to herd immunity.

### Influenza C evolution

Neither of the two participants who were identified as influenza C-positive by PCR generated sufficient deep sequencing reads for complete genomes to be assembled. Our deep sequencing of the nasopharyngeal swabs of both of our PCR-positive participants, produced much fuller genome sequence results for other RNA viruses apart from influenza C, as well as sequences from a range of bacterial species (Atkinson *et al* in preparation). We therefore do not think that the difficulty in detecting influenza C, or in generating complete genomes, is due to RNA degradation or other technical failure, but rather a true reflection of the rarity of the virus and a low viral titre in infected individuals.

In the individual with the 4 partial genome segment sequences, it is evident that C/Lancaster/1/2015 is a reassortant that does not fall into any of the genome constellations previously classified [Table 1 and Gatherer, 2010]. It contains a rare NS1/NS2 segment of the C/Sapporo/71 clade, related to sequences that were last observed in the late 1970s. Influenza C genomes sequenced since 2010 all have the C/Shizuoka/79 clade in the NS1/NS2 segment (Figure 5). C/Lancaster/1/2015 also has a rare NP segment of the C/Miyagi/1/93 clade, related to sequences that were last observed around 2000 (Figure 4) and typical of genome constellation 4a (Table 1). The other segments are within clades found more recently, although C/Lancaster/1/2015’s position within these clades is never close to any of the recent genome sequences (Supplementary Figures 1 and 2). The exact position of C/Lancaster/1/2015 on each segment’s phylogenetic tree is rarely well supported by Bayesian phylogenetics posterior probability density, but its location within each of the broader clades is well supported (see Supplementary Data Pack). We therefore conclude that its apparent reassortant nature is unlikely to be simply an artefact of partial sequence information.

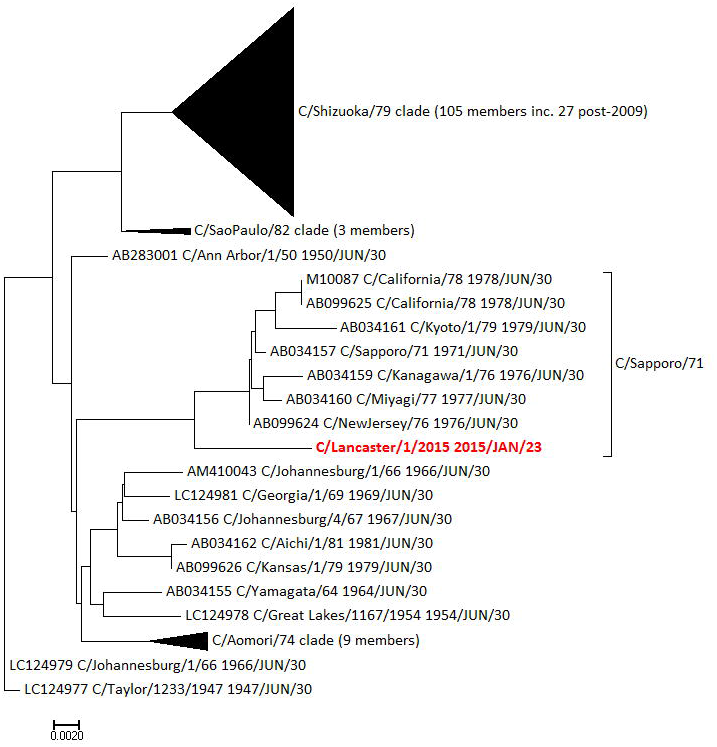

Tentative reconstruction of the reassortment event may be attempted. Gatherer [2010] defines genome constellation 4a as consisting of C/Sapporo/71, C/Miyagi/1/93, C/Sapporo/71 and C/Shizuoka/79 in segments 1, 5, 6 and 7 respectively. The corresponding clades for C/Lancaster/1/2015 are C/Sapporo/71, C/Miyagi/1/93, C/Sapporo/71 and C/Sapporo/71 (Table 1), suggesting that a strain of constellation 4a reassorted with one containing a C/Sapporo/71-clade segment 7. Since no strain containing a segment 7 of this clade has been seen since the 1970s and constellation 4a was only seen in the 1990s, it seems likely that the reassortment event occurred in the 1990s. This would also explain the dissimilarity of C/Lancaster/1/2015 in all of its segments, to other recently sequenced strains. We are tempted to speculate that this reassortant occurred locally in Lancaster, but in the absence of any other British genomes since C/England/892/1983 [Matsuzaki et al., 2016], which is itself incomplete, it is impossible to come a conclusion.

If this scenario is common in small isolated populations, influenza C diversity in terms of shifting genome constellations may be even greater than suggested from the available genomes. Rare strains may persist at low levels in small urban/rural locations, such as Lancaster. Our detection rate, at 0%, 0.7% or 1.4% depending on whether both samples, or merely one sample, is scored as positive, or whether asymptomatic individuals are included, is broadly similar to the 0.2% (frequency per symptomatic individual) found in another recently sampled British population [Smith et al., 2016]. The M1/CM2 (Figure 3) and NP segments (Supplementary Figure 3) for C/Lancaster/1/2015 have lower root-to-tip distances than expected under the assumption of molecular clock-like evolution. When this method is used on database-derived sequences, it is often taken as indicative of incorrect dating. However, given that we know precisely when our samples were collected, it is more likely to reflect a genuinely slower rate of evolution in these samples. The M1/CM2 segment of C/Lancaster/1/2015 is positioned in the phylogenetic tree near segments from the 1980s (Supplementary Figure 1) and the NP segment near segments from the 1990s and 2000 (Figure 4). This same phenomenon of slowed molecular clock, and aberrant positioning with the phylogenetic tree, has been seen in some strains of Zaire ebolavirus [Lam et al., 2015] and also in the 1977 “Russian Flu” H1N1 outbreak [Wertheim, 2010], and is thought be a consequence of the virus entering a host population where the serial interval – the time between infection of one host and the next in a transmission chain – is reduced and the virus therefore spends longer in a non-replicative state. For ebolavirus, this is assumed to be a non-typical animal reservoir host, and for Russian Flu possibly a laboratory freezer. Neither of these options would seem to be possible for influenza C, so it may simply be a cumulative result of low transmission rates within relatively small populations slightly delaying the average serial interval, conditions which could apply in Lancaster.

### Implications for vaccination strategy

We began this study with the premise that influenza C might be a candidate for inclusion in the seasonal influenza vaccine. Our results do not provide any support for the proposition that vaccination of adults is appropriate, a conclusion also reached by Smith et al. [2016]. Although we recruited 71 symptomatic individuals with a range of cold/flu-like symptoms, none of these was influenza C-positive, and none of the respiratory disease burden in Lancaster during our study period can be attributed to influenza C.

There may still be a case for vaccination of children in the light of published reports of serious respiratory disease caused by influenza C in that age group. [Calvo et al., 2006; Gouarin et al., 2008; Laxdal et al., 1966; Matsuzaki et al., 2007; Matsuzaki et al., 2006; Matsuzaki et al., 2003; Moriuchi et al., 1991; Peng et al., 1996; Principi et al., 2012; Shimizu et al., 2015; Takayanagi et al., 2009]. We recruited 6 participants in the <9 years age group but none were consented to allow serum sampling. In the single participant in the 10-19 year age group, anti-influenza C IgG levels were at <1 mg/dl and this individual is classified as seronegative (Figure 1). Whether the higher levels of anti-influenza C IgG found in our adults are as a consequence of a single exposure, or limited number of exposures, during childhood, or are maintained by recurrent possibly sub-clinical infections (as in our 2 positive participants) in adulthood, remains a matter of debate. The apparent slowing of evolutionary rate in the M1/CM2 and NP segments of C/Lancaster/1/2015, if caused by reduced average serial interval due to reduced number of infections in small isolated populations, possibly suggests the former.

**Table 1:** **Clade membership of segments of C/Lancaster/1/2015** based on Bayesian phylogenetic analysis and the prior clade and genome constellation classifications of Gatherer [2010]. The rightmost column lists those clades found in other segments sequenced from 2010 onwards. Segments 1 and 6 of C/Lancaster/1/2015 are outliers within clades found in other recent genomes, but segments 5 and 7 are not.

| Genome segment (encoded protein) | Clade of segment in Lancaster consensus, as defined by Gatherer [2010] | Genome constellation in which that clade is present | Clade(s) of other post 2009 genomes |
| --- | --- | --- | --- |
| 1 (PB2) | C/Sapporo/71 | All, except 5 | C/Greece/79;<br>C/Sapporo/71 |
| 5 (NP) | C/Miyagi/1/93 | 4a | C/pig/115/Beijing/81;<br>C/Greece/79 |
| 6 (M1/CM2) | C/Sapporo/71 | All, except 2 & 3 | C/Sapporo/71 |
| 7 (NS1/NS2) | C/Sapporo/71 | None: clade not seen since 1970s | C/Shizuoka/79 |

---

**Supplementary Figure 1: Neighbour joining tree rooted on C/Taylor/1233/1947 for segment 6** (M1/CM2), annotated with clades derived from Gatherer [2010] and confirmed by BEAST analysis (see Supplementary Data Pack), demonstrating the closer relationship of C/Lancaster/1/2015 (red) to M1/CM2 segments of the C/Sapporo/71 clade from the 1980s than to recent isolates. Scale: substitutions per site.

**Supplementary Figure 2: Neighbour joining tree rooted on C/Taylor/1233/1947 for segment 1** (PB2), annotated with clades derived from Gatherer [2010] and confirmed by BEAST analysis (see Supplementary Data Pack), demonstrating the closer relationship of C/Lancaster/1/2015 (red) to PB2 segments of the C/Sapporo/71 clade from the 1970s and 1980s than to recent isolates. Scale: substitutions per site.

**Supplementary Figure 3: Root-to-tip distance in a neighbour joining tree for segment 5** (encoding NP) of the influenza C genome. 96 full-length or near full-length genome segments (1809 bases) are used plus the 397 discontinuous bases of segment 5 derived from deep sequencing. C/Lancaster/1/2015 is less divergent from the root than it should be given its known sampling date, consistent with a perturbation of molecular clock-like behaviour in its lineage.

**Supplementary Figure 4: Root-to-tip distance in a neighbour joining tree for segment 7** (encoding NS1/NS2) segment of the influenza C genome. 134 full-length or near full-length genome segments (935 bases) are used plus the 288 discontinuous bases of segment 7 derived from deep sequencing. C/Lancaster/1/2015 has a degree of divergence from the root consistent with molecular clock-like behaviour in its lineage.

## Acknowledgments

KVA received a Service Increment from Teaching (SIFT) studentship from University Hospitals of Morecambe Bay (UHMB) National Health Service (NHS) Foundation Trust, UK and performed this work as part of the requirements for the degree of Master of Science (MSc). Rosetrees Trust, UK, provided additional funding for deep sequencing (grant M395).

## Ethics Statement

Informed consent was obtained from adult volunteers and supported assent from juveniles with prior informed parental consent. Ethical approval was granted by the UK National Research Ethics Service (NRES), reference 14/LO/1634, Integrated Research Application System (IRAS) Project 147631. The project was registered with the National Institute of Health Research (NIHR), UK as part of the NIHR Clinical Research Network (UKCRN) Portfolio, ID 17799.

## Data Accessibility Statement

The Supplementary Data Pack containing statistical analyses on volunteers and ELISAs, BAM files and reference genomes for genome assemblies, genome fragments too short for inclusion in GenBank, BEAST inputs and outputs, TempEst inputs and outputs and pipeline Perl scripts, are available from: doi://10.17635/lancaster/researchdata/111

